# MyoNet and AlveoliNet: A hybrid 3D neural-net pipeline for the instance segmentation and scoring of epithelial cells in intact mammary alveoli

**DOI:** 10.64898/2026.05.26.727878

**Authors:** RF Monks Colin, A Fester Sarah, Cassidy J Nicks, Kiarra Coger, Yukiko U Inoue, Takayoshi Inoue, Jenifer Monks

## Abstract

Recent advances in tissue clearing protocols such as DISCO, CUBIC, Clarity, FUnGI, and PEGASOS have revolutionized our ability to label and image intact 3-dimensional (3D) biological structures using fluorescence microscopy. The lactating mammary gland particularly benefits from clearing due to its high degree of tissue opacity. Cleared mammary gland images are strikingly beautiful and complicated but are difficult to fully interpret without developing a series of quantitative techniques and assays to analyze and compare them. These approaches will ultimately be as varied as the biology each scientist wishes to study. Here, we present one strategy based on a modular, hybrid, deep-learning approach and classical image processing that can segment and measure alveoli, cell nuclei and myoepithelial cells in intact mammary tissue. We have developed two original, three-dimensional (3D) U-shaped encoder-decoder networks (U-Nets), AlveoliNet and MyoNet, and combined these with CellPose3 nuclear instance segmentation and SlideBook/SlideBook Synergy binary mask operations. This approach can be used to easily score 100,000s of cells in intact tissue and differentiated glands at different developmental stages, genetic backgrounds, or treatments. We demonstrate the utility of this approach for quantifying the change in proportion of myoepithelial cells over the pregnancy-lactation transition, driven by endoreplication in the gland postpartum. We present a complete methods pipeline for other laboratories to utilize our approach in their own studies using standard desktop computers.

## Introduction

Tissue staining and clearing techniques to observe glandular development of the mouse mammary gland were developed over 75 years ago, using Hematoxylin and Carmine alum dyes and organic solvents. (protocols given in ^1,2^, respectively). These protocols have been used by the field to carefully elucidate the hormonal and molecular signaling pathways that contribute to ductal growth and branching during puberty, changes in the gland during estrus cycling, glandular development during pregnancy, regression during involution, and stages of cancer progression.^3–5^ However, during lactation, the density of the mammary gland increases due to increased milk-secreting mammary alveoli, depletion of the adipocyte fat depots, and the presence of highly opaque milk. Imaging of the mammary gland at depth during lactation suffers from increased scatter and shadowing by overlaying structures. Thus, traditional methods of scoring histological, developmental, or pathological differences between mammary glands in genetically modified mice have relied on analysis of tissue sections. Using standard histological stains or immunolabeling, a trained histopathologist may score the tissue sections to create image analysis matrices.^6^ Traditional histological scoring requires a trained histologist, is extremely time-consuming for large tissue cohorts, and generates only semi-quantitative data. Alternatively, 2-dimensional (2D) images may be segmented by software for quantitative analysis.^7^ However, analysis of tissue sections has significant limitations such as the inability to reconstruct the 3D morphology of the tissue.

The intact mammary gland is a geometrically complex, cellularly heterogeneous structure. The two alveolar epithelial cell types present during lactation, primary milk-secreting luminal epithelial cells and contractile myoepithelial cells, are physically in proximity and are not easily disambiguated at light microscope resolution. Additionally, the tissue is highly heterogeneous, containing regions of alveoli, adipose tissue, vasculature, and inflammatory cells.^8^ Recently, mammary gland biologists have begun to utilize clearing techniques that enable the imaging of genetically encoded fluorescent proteins, as well as cells/structures stained with vital dyes or antibodies.^8–12^ Improvements to confocal and multiphoton microscopy now enable imaging deeper into the tissue. Additionally, computational method development and image processing power allow for the capture of large, 3D datasets, visualizing the structures of the mammary gland at ever-increasing resolution.^8,13^ What is needed is a coherent method for analysis of the lactating gland that can be shared between groups, to elucidate the comprehensive changes in the gland over development, or with genetic or pharmacologic manipulation. To this end, we utilized and developed a series of deep-learning approaches to segment and analyze cleared, intact tissue for nuclei and total myoepithelial count.

To count the total nuclei in a 3D section of gland, we stained the tissue using the far-red dye RedDot2 and used recently developed and improved instance segmentation methods. We were successful with both the B-box-first approach^14–16^ in 3D using Stardist, and the flow-field approach ^17^ in 3D using CellPose, for scoring nuclei in cleared and stained tissue. Ultimately, CellPose gave the most consistent and accurate nuclear counts for our data.

Traditionally, semantic segmentation strategies have been effectively employed for delineating the boundaries of bulk structures in 3D.^18^ To score nuclei that reside within an alveolar structure, we created and trained a specialized 3D U-Net that we have named AlveoliNet. AlveoliNet takes as input 3D images of the nuclear stain RedDot2 at low resolution (30 microns x 30 microns x 30 microns). RedDot2 clearly stains DNA and has the additional benefit of yielding some diffuse staining in alveoli but not other cell types in the mammary gland. AlveoliNet was built using a training set created from hand-edited, intensity-segmented, 3D images of alveoli. Its output is a clean delineation of voxels that are within a 3D alveoli. The network has been made publicly available for others in the community to use.

The contractile, myoepithelial cells in the lactating mammary gland are impossible to study in 2D sections but are readily visible in 3D projections. (**Fig. 1**) To segment myoepithelial cells, we have designed and trained a specialized U-Net to segment the myoepithelial voxels in our 3D data. We have named this network MyoNet. This network was trained using augmented hand-delineated examples and carefully intensity-segmented examples across two separate labeling strategies. The network has also been made publicly available for others in the community to use.

**Figure 1:**
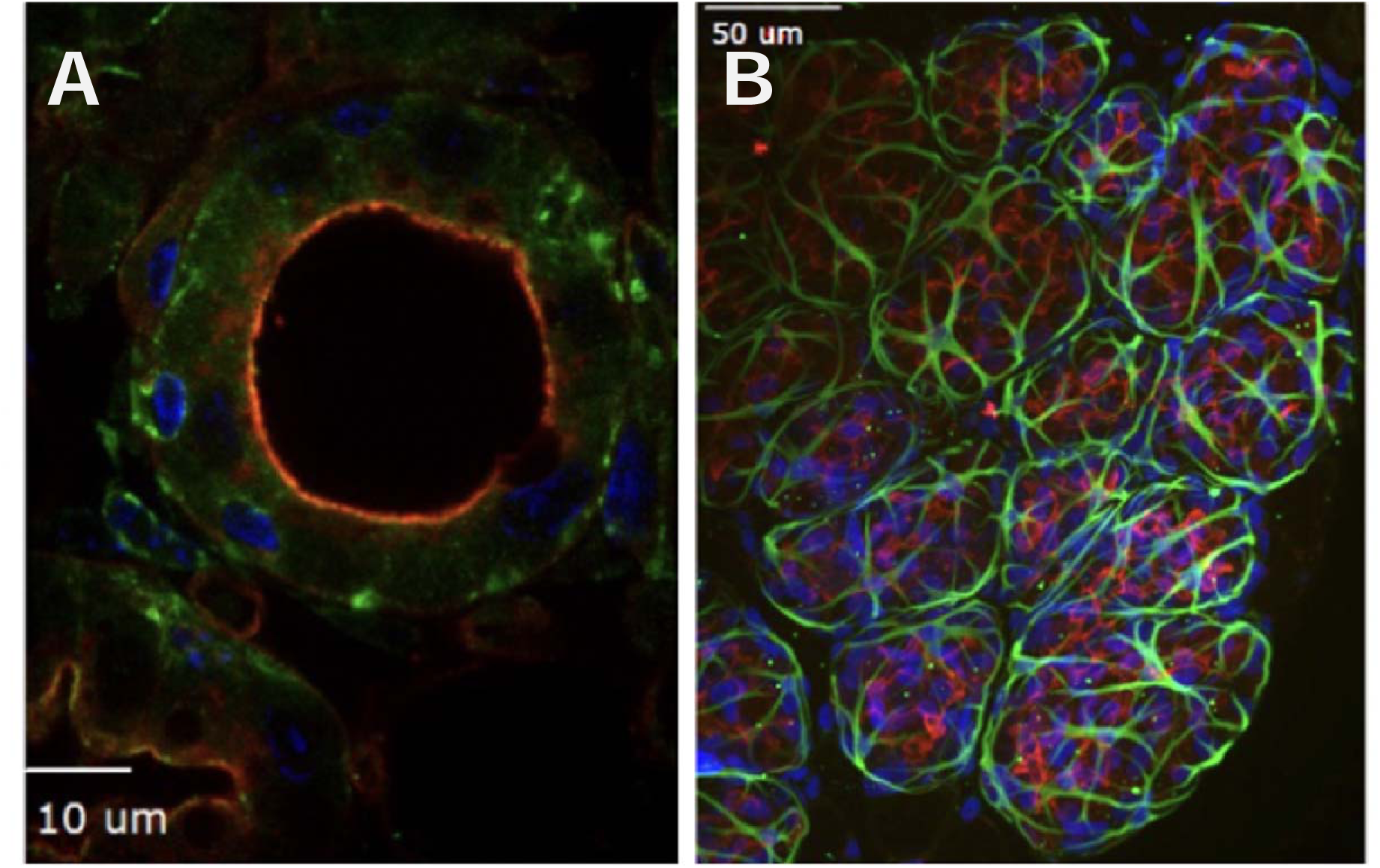
Myoepithelial cells must be imaged and characterized in 3-dimensional tissue. Myoepithelial cells visualized with smooth muscle actin immunostaining (green), co-stained with DAPI (blue) to visualize nuclei and wheat germ agglutinin (red) to label the apical plasma membrane of the secretory epithelial cells. **A.** A single-plane tissue section, **B**. a 3D maximum intensity projection view through a 50 micron-thick tissue.

Finally, to determine the fraction of total cells that are myoepithelial cells, we employed classical 3D image processing, such as object-level logical mask AND operations and size gating to eliminate very small objects.^19^ These operations were performed in SlideBook and a sidecar application that uses the Synergy module protocol to communicate with SlideBook.

The final analysis represents a hybrid approach that employs both the semantic segmentation of custom U-Nets and the flow-field approach (CellPose) to create objects that can be logically interacted with and scored using classical mask operation. (**Table 1**) Tens of thousands of cells were ultimately scored in intact glands using this approach within hours.

**Table 1:** Comparison Summary: Segmentation Strategies.

| Model/Strategy | Segmentation Type | Primary Strength | Best Use Case |
| --- | --- | --- | --- |
| Custom U-Net | Semantic (or specialized 3D Instance) | Optimized for **specific, proprietary 3D structures** due to custom training. | **High-precision segmentation** of rare or non-standard 3D structures. |
| CellPose | Flow-Field (Instance) | **Generalization and Speed** (no training required for standard cells/nuclei). | Rapid, robust segmentation and counting of individual nuclei and cells in 2D or 3D images. |
| U-Net + CellPose | Hybrid (Semantic + Instance) | **Precision in Classification and Counting** | Phenotyping cell populations in crowded tissue where different structures define cell identity. |

## Methods

### Animal studies

Mice were housed in the University of Colorado Denver Anschutz Medical Campus, AAALAC-accredited (#00235, Approved 07/11/2024), Center for Comparative Medicine. Mice had *ad libitum* access to food (Harlan 2920x), in micro-isolator cages with automated air and water, on a 10:14 h dark/light cycle, maintained at 72°F. All animals were provided with cotton nesting material enrichment, and breeders and experimental females were additionally provided with shredded paper to construct an enclosed nest. Experimental females were placed with proven C57Bl/6/J stud males. The presence of a vaginal plug was recorded and counted as pregnancy day 1 (P1). Dams were individually housed before parturition. The day a litter was first seen was counted as lactation day 1 (L1). Dams were harvested on P19 or L4 by CO_2,_ followed by intracardiac perfusion of Phosphate-Buffered Saline (PBS), followed by 2% then 4% paraformaldehyde. Freshly collected mammary glands (thoracic) were further fixed in 4% paraformaldehyde in PBS at room temperature overnight, then moved into PBS-azide at 4°C for long-term storage. All procedures were approved by the Institutional Animal Care and Use Committee of the University of Colorado Denver Anschutz Medical Campus (Monks, protocol #1200).

Oxtr-1xPA-tdTomato mice (JAX 037580, RIKEN RBRC12289)^20^ were a kind gift of Dr. Yukiko U. Inoue, National Institute of Neuroscience, Tokyo, Japan. These mice were backcrossed to C57Bl/6/J mice for four generations before experiments were started. All the glands used to create the data presented in this manuscript are from heterozygous dams, which we validated did not have any reproductive deficiency. (data not shown, available upon request). tdTomato expressing glands were co-stained with RedDot2 (1:200, ThermoScientific, NC0772106,) overnight at 37°C and then washed with PBS-azide. Clearing was performed using a modified CUBIC protocol (TCI: CUBIC-L #T3740, followed by CUBIC-R(+M) #T3741).

For immunostaining, 1 cubic mm pieces were cut from the middle of the gland and, freely floating pieces were immunofluorescently labeled with Rabbit anti-Smooth Muscle Actin (SMA, 1:5000, Abcam ab5694) and Donkey anti-Rabbit antibody-Alexa 555+ conjugated secondary antibody (Invitrogen A32794), co-stained with RedDot2, and cleared using a modified iDISCO protocol.^21^ Note: tdTomato fluorescence does not survive clearing with iDISCO reagents.

## Microscope hardware/software and Imaging conditions

Three-Dimensional Confocal images were collected on a Marianas SDC microscope (Intelligent Imaging Innovations, Inc., Denver, CO, USA) using a Zeiss 40x 1.3 NA Objective (Carl Zeiss, GmbH, Jena, Germany) and a 50-micron pinhole CSUW spinning disk confocal (Yokogawa Electric, Kanazawa, Japan). The images were captured in 16-bit using SlideBook 2025 software (Intelligent Imaging Innovations, Inc., Denver, CO, USA), a Prime95B scientific CMOS camera (Teledyne Photometrics, Tucson, AZ, USA), and appropriate emission filters (Chroma Technology, Bellows Falls, VT, USA). Data were collected as 3D image stacks with 0.3 µm spacing between planes. Stack depths were constrained by the objective lens working distance (350 microns) and the geometry of individual cleared mammary glands. Image stacks were between 100 and 130 µm in depth (Z). The tdTomato and Alexa555 channels were collected using the 100 mW 561 laser line from a LaserStack v4 (Intelligent Imaging Innovations, Inc., Denver, CO, USA). RedDot2 was collected using a 150 mW 640 laser line. Exposure times were optimized per sample but ranged between 500 ms and 1000 ms per channel. 3D image frames were 1200x1200 pixels. Image stacks were between 300 and 400 planes. The resulting 2-channel 3D images ranged between 2.0 and 2.5 GB total.

The original SlideBook (.sldy) files are available https://figshare.com/s/518e4fd1a86d21801117 and can be opened in SlideBook, SlideBook Reader (free), FIJI, or any program supporting BioFormats.^22^

## Semantic Network Creation (MyoNet)

Initial training masks were produced by the manual delineation of myoepithelial cells using the virtual reality software syGlass (Morgantown, WV, USA). Training sets were created from 3D image stacks (1200x1200x600) of tdTomato-labeled myoepithelial cells. Adaptive Histogram Equalization (AHE) was applied to the training data, and the resultant output was added as an additional reference channel. This AHE channel was used as a guide to help the annotator differentiate between faint myoepithelial cells and the background. The AHE channel was not utilized during the training process of the U-Net. The entirety of the myoepithelial cell was masked using the masking tool in syGlass. (**Movie 1**) Masks were exported with 3D labeling for future U-Net training (MyoNet). Masks and data were downsampled using the SciPy zoom function^23^ with a scale factor of 0.35. Once downsampled, the images were patched into training data using the library patchify.^24^ There were four total images used to assemble a training dataset. A total of 3523 patches were produced: 1835 patches from myo1 using a stride length of 4, 430 patches from myo 2 (m2c1c4) using a stride length of 10, 238 patches from myo (3m3c4c1) using a stride length of 10, and 1020 patches from myo 4 (m4c1c7) using a stride length of 10. All patches were sized 32 x 32 x 32 voxels. Any patches produced that had no masked areas were discarded. This was done to help control the class imbalance between foreground masked regions and background unmasked regions. The data was split, with 70% of the images being used for training data and 30% being used for validation data.

The Python library, Volumentations, was used to apply augmentations to the training data.^25^ This was done to extend the training set and reduce the likelihood of the U-Net overfitting the training data. Prior to the augmentation being applied, the image was normalized by the maximum value within the image. The augmentations selected were elastic transformation, random gamma adjustment, random Gaussian noise addition, random 90-degree rotations, and random flips along each axis of the image. Each augmentation was applied randomly with different chances of occurrence. Elastic transformation, random gamma adjustment, and random Gaussian noise addition had a 40% possibility of being applied to an image. Random 90-degree rotations and random flips along each axis of the image had a 50% chance of being applied to the image. After the augmentations were applied, the values within the images were clipped to fall between zero and one. The augmentation pipeline was active for each image. No augmentations were applied to images in the validation set.

The U-Net (MyoNet) was trained using the Alpine system (a supercomputer with 80 NVIDIA A100 GPUs) at the Boulder Supercomputing Center for 100 epochs with a batch size of four. The Adaptive Moment Estimation (Adam) optimizer was used, and the learning rate was set to 0.001.^26^ The loss function used was Tversky Loss, with the alpha and beta parameters set to 0.2 and 0.8, respectively.^27^ This was done to penalize false negatives during training. Network weight selection was made with respect to the lowest validation loss. The validation loss was also Tversky Loss, using the same alpha and beta parameters as the training loss settings. Epoch 99 had the lowest validation loss with a value of 0.0998 and a validation accuracy of 0.98. The Network was saved as a PyTorch weights file^28^ for use in CellNet 2.0 (Intelligent Imaging Innovations, Inc, Denver, CO). This weights file is publicly available for general use at FigShare. (https://figshare.com/s/518e4fd1a86d21801117)

CellNet 2.0 is a software platform developed by Intelligent Imaging Innovations (3i) for the generation of training datasets, data review, augmentation, neural network training, and model inference. The application utilizes 3i Synergy technology to provide direct access to SlideBook image data and generation of network-derived segmentation masks. CellNet 2.0 training and inference are optimized for CUDA acceleration on NVIDIA GPUs, but inference can also be performed on Intel CPUs using OpenVINO (Intel Corporation, Santa Clara, CA, USA). CellNet 2.0 supports 2D and 3D U-Net architectures implemented using TensorFlow^29^, PyTorch,^28^ and ONNX-based model deployment.^30^

MyoNet was applied, via CellNet 2.0, producing SlideBook masks that define the voxels of the 3D myoepithelial cells. Masks are saved with the original data in SlideBook and are available with the data online. (https://monkslab.org/portal/)

## MyoNet Revisions and Fine Tuning

The initial MyoNet produced accurate segmentation on tdTomato-stained samples but tended to under segment samples labeled with smooth muscle actin (SMA). To make a more generally useful (less brittle) network, MyoNet was fine-tuned with a selection of 3000 32x32x32 intensity-segmented SMA data cubes. This training set was downsampled and augmented as described above. The network was re-run for an additional 100 epochs. The training set and masks are available. (https://monkslab.org/portal/) This fine-tuned network is the final MyoNet and was used to segment all the data reported here.

## AlveoliNet Creation, Revisions, and Fine Tuning

Initial training masks for an additional U-Net to delineate alveolar structures from stroma (AlveoliNet), were produced by intensity segmentation, which were hand edited using SlideBook 2025 and Synergy Conductor (Intelligent Imaging Innovations, Inc., Denver, CO, USA). Training sets were created from 10 3D image stacks (1200x1200x600) of RedDot2 labeled mammary tissue. Masks were exported with 3D labelings for future U-Net training (AlveoliNet). Masks and data were downsampled using the training set creation functions of CellNet 2.0 using a scale factor of 0.1. Once downsampled, the images were patched into 3000 separate 40x40x40 training data patches. Patches deliberately included negative examples where no alveoli exist. The training set and masks are available. (https://monkslab.org/portal/)The data was split, with 70% of the images being used for training data and 30% being used for validation data.

The 3D U-Net network was trained in CellNet 2.0. CellNet 2.0 provided augmentations to the training data that adjusted scale, signal-to-noise, rotation, geometric distortion, and photobleaching. This was done to extend the training set and reduce the likelihood of the U-Net overfitting the training data. Prior to the augmentation being applied, the image was normalized by the maximum value within the image. Each augmentation was applied randomly with different chances of occurrence. Elastic transformation, random gamma adjustment, and random Gaussian noise addition had a 40% possibility of being applied to an image. Random rotations and random flips along each axis of the image had a 50% chance of being applied to the image. After the augmentations were applied, the values within the images were clipped to fall between zero and one. The augmentation pipeline was active for each image. No augmentations were applied to images in the validation set.

The 3D U-Net (AlveoliNet) was trained on a Dell Precision 7960 Tower workstation (Dell Technologies, Round Rock, TX, USA) equipped with an NVIDIA RTX A4500 GPU (NVIDIA Corporation, Santa Clara, CA, USA) running Windows 11 Pro (Microsoft Corporation, Redmond, WA, USA) for 120 epochs using a batch size of four. The Adaptive Moment Estimation (Adam) optimizer was used with a learning rate of 0.001. The loss function was Tversky loss, with α = 0.2 and β = 0.8 to preferentially penalize false negatives during training. Network weights were selected based on the lowest validation loss. Validation loss was also computed using Tversky loss with identical α and β parameters. Epoch 108 yielded the lowest validation loss (0.0999) with a voxel-wise validation accuracy of 0.91. AlveoliNet was subsequently fine-tuned on two independently edited segmentation datasets for an additional 100 epochs to improve consistency across datasets. The final AlveoliNet model was saved as a PyTorch weights file (.pth) for use in CellNet 2.0 (Intelligent Imaging Innovations, Denver, CO, USA). The weights file is publicly available at Figshare.(https://figshare.com/s/518e4fd1a86d21801117)

Like MyoNet, AlveoliNet was applied via CellNet 2.0 to produce SlideBook masks that define the location of the voxels that are the 3D Alveolar region. Masks are saved with the original data in SlideBook and are available with the data online. (https://monkslab.org/portal/)

## Running the analysis/implementing the U-NETs and CellPose

Nuclei were segmented using Cellpose 3 executed within CellNet 2.0. The pretrained nuclei model was used with default inference parameters, including automatic nuclear diameter estimation by Cellpose. Segmentation was performed in 3D mode using Cellpose-recommended normalization settings and default probability and flow thresholds. Inference was performed using the same hardware configuration described for AlveoliNet training. Segmentation results were exported as instance masks, in which each nucleus was assigned a unique label, and subsequently imported into SlideBook 2025 for downstream analysis.

## Validation

All masks were evaluated in each data set by empirical simple visual evaluation prior to proceeding to the next step.

## Finalizing the analysis in SlideBook

Instance-segmented nuclei generated by Cellpose 3 within CellNet 2.0 were filtered using object-level volumetric intersection analysis in SlideBook 2025 and Synergy Conductor. Nuclear objects were intersected with the binary segmentation mask generated by AlveoliNet. Nuclei exhibiting greater than 80% volumetric inclusion within the AlveoliNet mask and volumes exceeding 1000 voxels were retained and classified as alveolar nuclei. The size threshold was applied to exclude biologically implausible small objects and segmentation artifacts.

Alveolar nuclei were subsequently intersected with the binary segmentation mask generated by MyoNet. Alveolar nuclei exhibiting greater than 50% volumetric inclusion within the MyoNet mask were classified as myoepithelial nuclei. All nuclei were retained as instance-segmented objects throughout the analysis to preserve object identity for downstream quantitative analysis.

## Statistics

Statistical analyses were performed using GraphPad Prism version 11 (GraphPad Software, Boston, MA, USA).

## Results

Image acquisition was performed as described in the Methods section. A training dataset as generated from 3D image volumes of lactating mouse mammary gland expressing tdTomato under control of the oxytocin receptor promoter, which is highly expressed in myoepithelial cells was targeted for segmentation. Figure 2 illustrates preparation of the training dataset, including examples of manually segmented ground-truth masks (**Fig. 2A**) and subdivision of image volumes into training patches (**Fig. 2B**). Training data consisted of 3523 volumetric image patches derived from manually segmented mammary gland datasets. Preparation of the training dataset is also demonstrated in Supplementary Movie 1.

**Figure 2:**
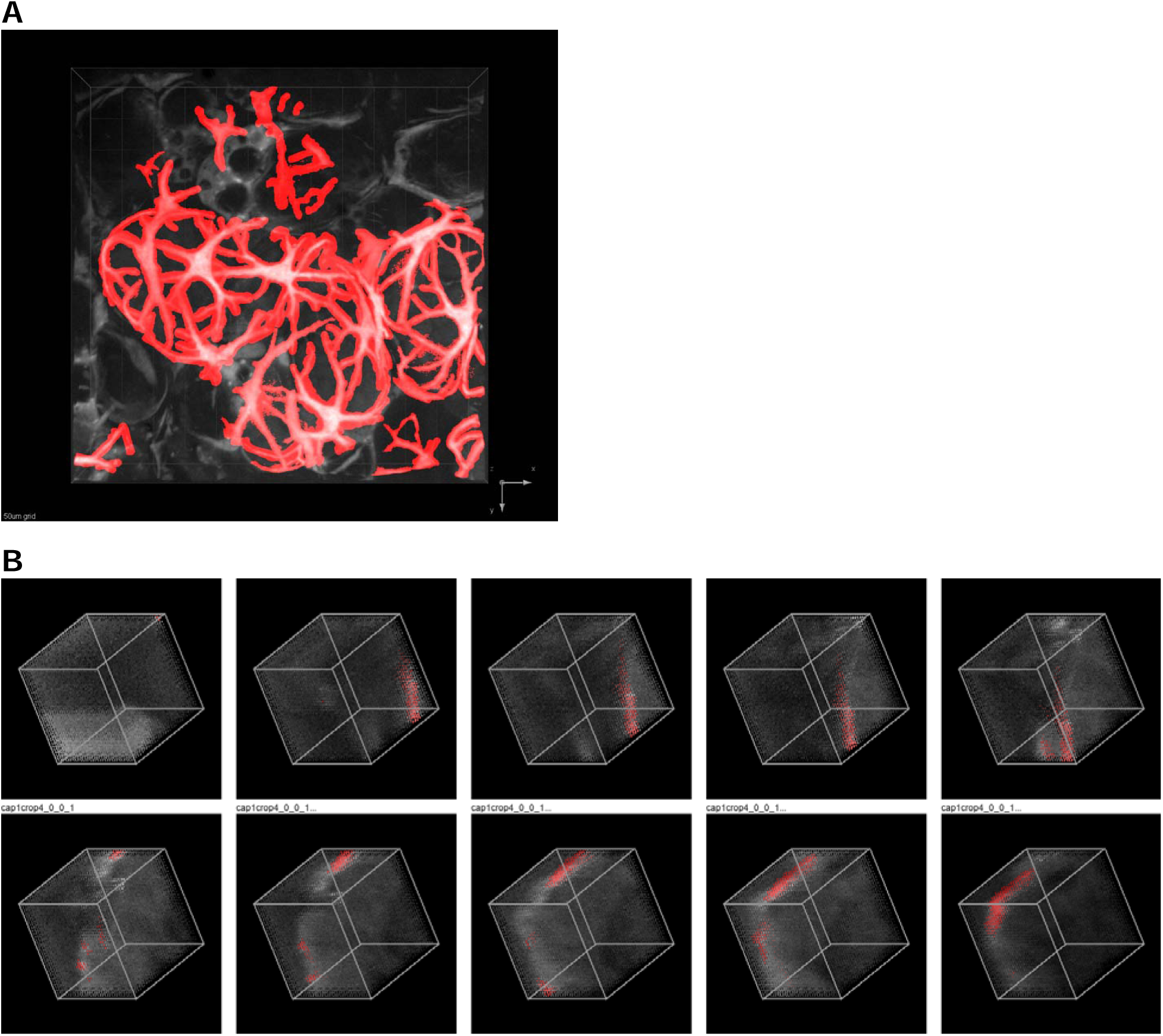
Training Set preparation. **A.** 3D rendering of tdTomato myoepithelial cells (black and white). 3D hand segmentation for training set creation is shown in red. Grid lines in rendering are 50 microns. (see also Movie 1) **B.** Image subdivision strategy for training data preparation. Whole images (1200×1200×600 voxels) were first divided into 8 equal 3D blocks. Selected blocks containing masks were downsampled (scale factor 0.35) and then patched into 32×32×32 voxel patches for MyoNet training.

Multiple network architectures and training strategies were evaluated during development prior to selection of the final MyoNet and AlveoliNet architectures. The final network architecture consisted of a fully convolutional 3D U-Net designed for volumetric semantic segmentation of myoepithelial structures within intact mammary gland tissue volumes. (**Fig. 3**) The network utilized a multiscale encoder-decoder structure with skip connections to preserve fine structural detail while incorporating broader volumetric context. This architecture was selected due to its ability to accurately segment thin and spatially complex myoepithelial processes across heterogeneous 3D tissue regions.

**Figure 3:**
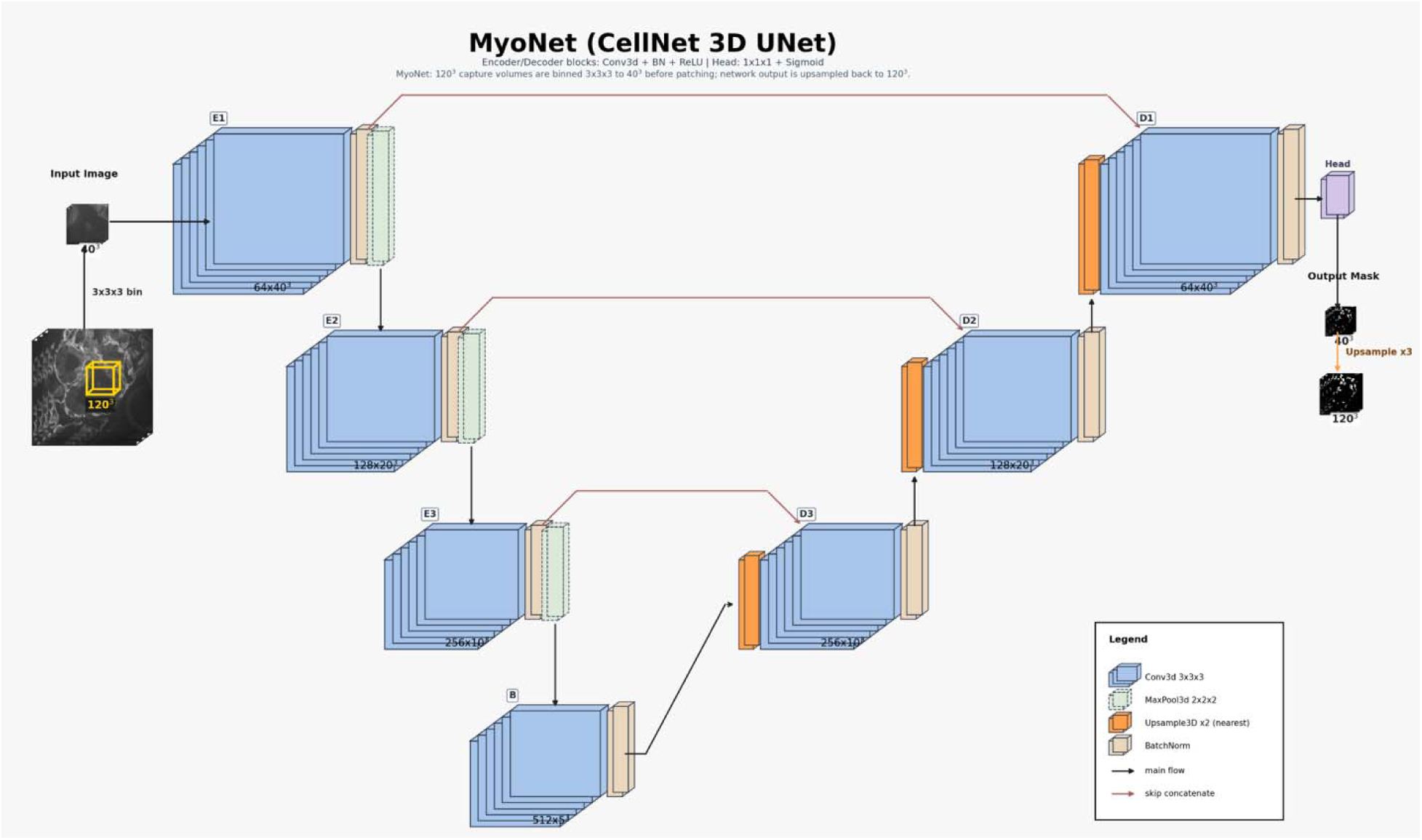
MyoNet (CellNet 3D U-Net) architecture. Capture volumes blocks of size 120³ are binned 3×3×3 to 40³ before network inference; the predicted mask is later upsampled ×3 back to 120³. The encoder (E1–E3) applies Conv3d 3×3×3 (+ BatchNorm + ReLU) followed by MaxPool3d 2×2×2, increasing channels while reducing spatial size (64×40³ → 128×20³ → 256×10³). The bottleneck (B) operates at 512×5³. The decoder (D3–D1) restores resolution with nearest-neighbor Upsample3D ×2 and Conv3d blocks (256×10³ → 128×20³ → 64×40³). Red skip connections concatenate encoder BatchNorm features into the matching decoder Conv3d stage. A 1×1×1 head with sigmoid produces the 40³ output mask. Blue blocks, Conv3d; tan, BatchNorm; green dashed, MaxPool3d; orange, Upsample3D; purple, head; black arrows, main flow; red arrows, skip concatenate.

Application of MyoNet to raw 3D mammary gland image volumes from pregnant and lactating tissue generated spatially precise volumetric segmentation masks (yellow, **Fig. 4C, D**) corresponding to tdTomato-positive myoepithelial cells. (red, **Fig. 4A, B**) MyoNet generated contiguous volumetric segmentations throughout intact tissue volumes and accurately segmented both strongly labeled and weakly labeled myoepithelial structures within dense alveolar tissue. The final fine-tuned MyoNet model also successfully generalized to smooth muscle actin (SMA)-labeled datasets (**Movie 2**), demonstrating robustness across labeling modalities.

**Figure 4:**
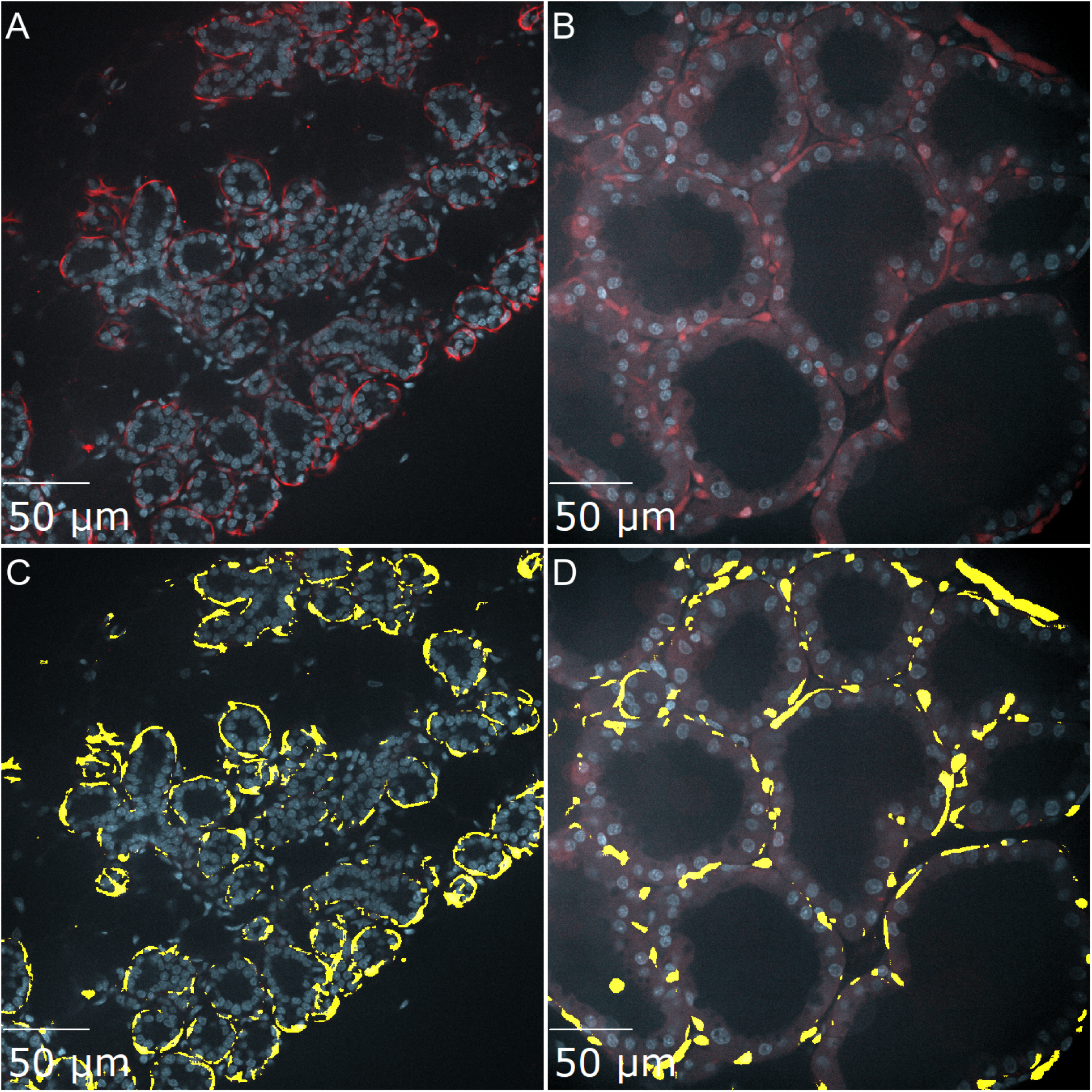
Results of MyoNet. Images captured at P19 (A,C) and L4 (B,D). A and B are 2D sections of 3D datasets showing DNA in blue and tdTomato myoepithelial cells in red. Figure C and D show the results of MyoNet segmentation in Yellow. Note that MyoNet provides a precision segmentation across the two distinct morphologies.

Nuclei were segmented using Cellpose 3 and combined with MyoNet and AlveoliNet segmentation outputs through object-level volumetric intersection analysis in SlideBook 2025 and Synergy Conductor. Integration of MyoNet, AlveoliNet, and Cellpose enabled automated object-level quantification of myoepithelial nuclei within large 3D mammary gland tissue datasets. Applying the MyoNet-AlveoliNet-CellPose segmentation pipeline to a single dataset results in the identification of all secretory mammary epithelial cells (sMEC) and all myoepithelial cells (myoEC) in the 3D image. (**Fig. 5**)

**Figure 5:**
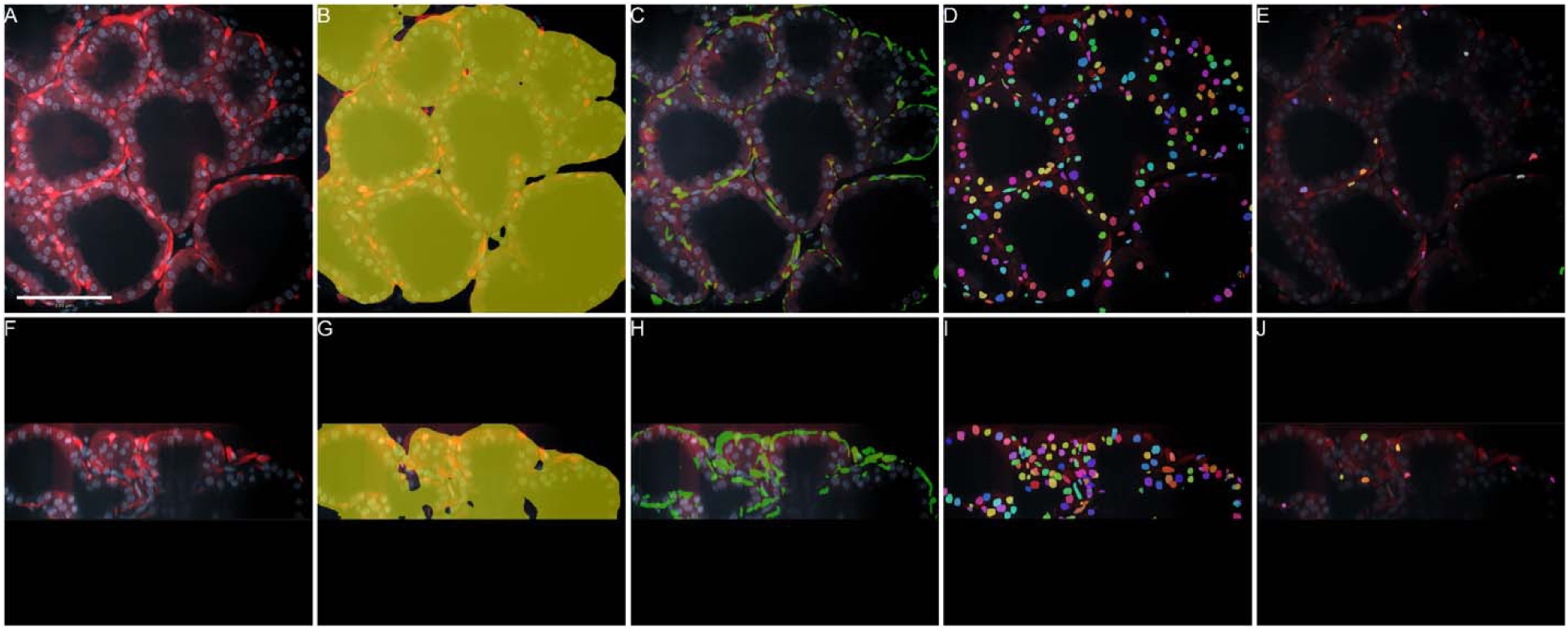
Results of MyoNet-AlveoliNet-CellPose segmentation pipeline. **A.** An XY slice of a 3D image of tdTomato labeled myoepithelial cells (Red) and RedDot2 labeled DNA (Blue) with a 100 µm scale bar shown. **F**. An orthogonal XZ slice of the same volume. **B.** and **G**. The same XY and XZ slices with the result of AlveoliNet overlayed in yellow. **C.** and **H.** show the same XY and XZ slices with the result of MyoNet overlayed in green. **D.** and **I**. show the same XY and XZ slices with the result of CellPose instance segmentation ANDed with AlveoliNet showing each alveolar nucleus overlayed in a rainbow pallet. **E.** and **J.** show the same XY and XZ slices with the results of the entire pipeline: myoepithelial nuclei overlayed in a rainbow pallet created by the AND of the result of MyoNet and the alveolar nuclei. These final panels (E and J) are the resultant myoepithelial nuclei.

The MyoNet-AlveoliNet-Cellpose analytical pipeline was subsequently applied to image datasets collected from four pregnancy day 19 (P19) animals and four lactation day 4 (L4) animals. Each developmental stage included a variable number of technical replicate image volumes acquired from individual mice. Datasets included tissues imaged for endogenous tdTomato fluorescence or immunostained for smooth muscle actin (SMA). Quantitative results for individual image volumes are summarized in Table 2, with aggregate results shown in Figure 6. Quantitative analyses revealed the expected change in the ratio of sMEC to myoEC, (**Fig. 6A**) alternatively shown as a decrease in the fraction of myoepithelial cells which occurs with endoreplication of the sMEC between P19 and L4. (**Fig. 6B**) Together, these results demonstrate that the combined MyoNet-AlveoliNet-Cellpose workflow enables quantitative identification of myoepithelial nuclei within intact 3D mammary gland.

**Figure 6:**
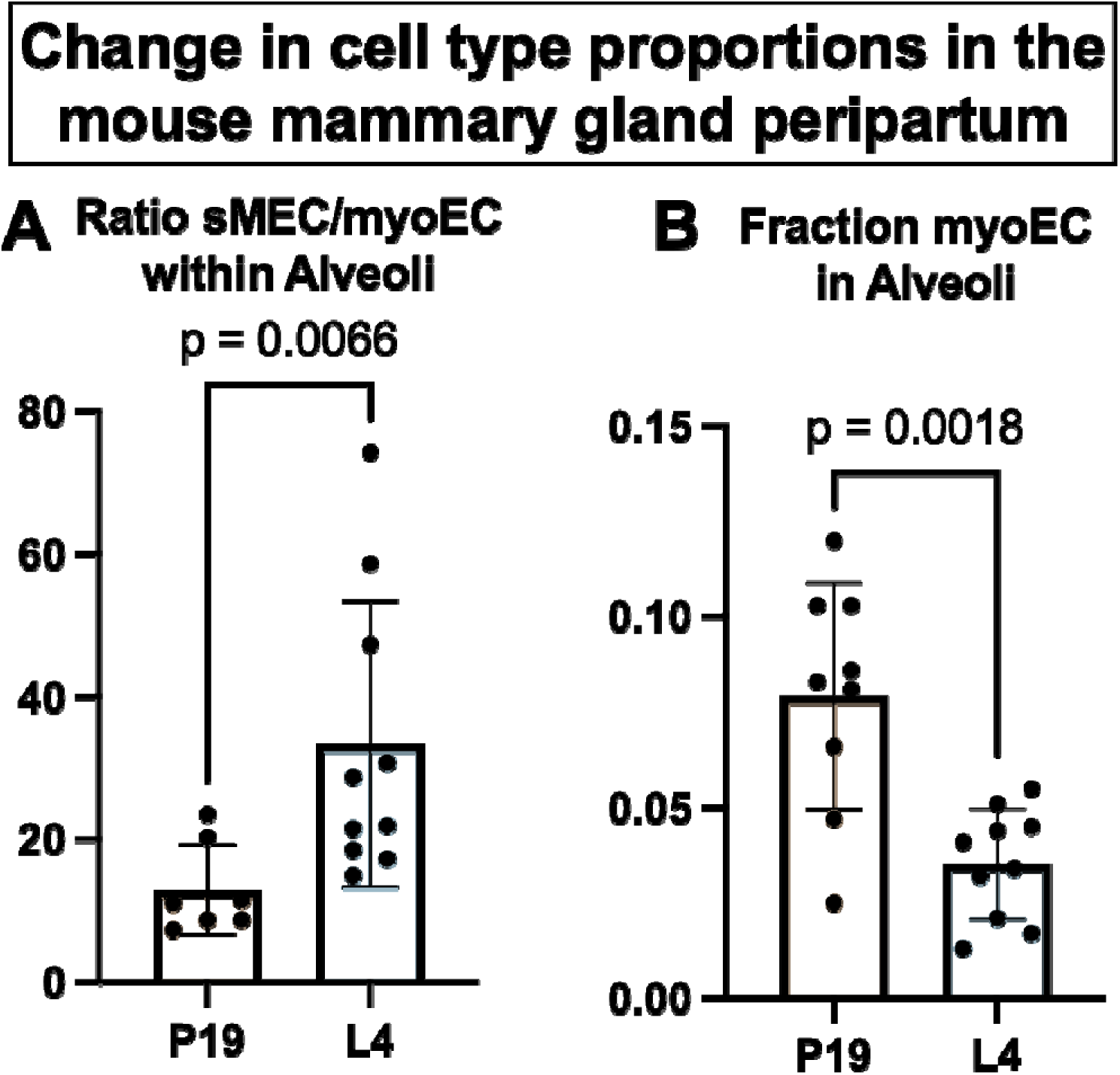
Change in cell type proportions in the mouse mammary gland peripartum as determined using the MyoNet-AlveoliNet-CellPose pipeline. A. Shown are the ratio of secretory mammary epithelial cells (sMEC) to myoepithelial cells (myoEC) within segmented alveoli. B. Shown are the fractions of myoEC in segmented alveoli. Each dot represents an individual data set in listed Table 2. All mice and data sets from each timepoint were included in aggregate data. For statistical analysis, a Mann-Whitney nonparametric test was used to compare ranks.

**Table 2:** Quantification of myoepithelial cells in mouse mammary gland tissue using the MyoNet-AlveoliNet-Cellpose analytical pipeline. Reported measurements include image dataset identifier, mouse identifier, developmental stage, total nuclei detected within the 3D tissue volume, total myoepithelial nuclei identified, and the fraction of nuclei classified as myoepithelial. Intermediate segmentation outputs from each stage of the analytical pipeline are also reported, including Cellpose nuclei, Cellpose AND MyoNet myoepithelial nuclei, Cellpose AND AlveoliNet alveolar nuclei, and Cellpose AND AlveoliNet AND MyoNet alveolar myoepithelial nuclei.

| File | Mouse ID | Stage | Nuclei | myoEC Nuclei | Alveolar Nuclei | Alveolar myoEC Nuclei |  | Alveolar ratio MEC/myoEC | Fraction myoEC in Alveolar mask | Fraction sMEC in total section | Fraction myoEC in total section | overestimated myoEC without AlveoliNet |
| --- | --- | --- | --- | --- | --- | --- | --- | --- | --- | --- | --- | --- |
| Condition 1 | 476879R L4 TOM+ | L4 | 10931 | 381 | 9507 | 197 |  | 47.3 | 0.021 | 0.85 | 0.035 | 1.93 |
|  |  | L4 | 9341 | 374 | 8419 | 266 |  | 30.7 | 0.032 | 0.87 | 0.04 | 1.41 |
| Condition 1_o | 476879R L4 TOM+ | L4 | 4423 | 129 | 3953 | 176 |  | 21.5 | 0.045 | 0.85 | 0.029 | 0.73 |
|  |  | L4 | 5452 | 170 | 5018 | 219 |  | 21.9 | 0.044 | 0.88 | 0.031 | 0.78 |
|  |  | L4 | 4880 | 313 | 4746 | 261 |  | 17.2 | 0.055 | 0.92 | 0.064 | 1.2 |
| Condition 2_o | 476879LL L4 TOM+ | L4 | 3407 | 160 | 3329 | 171 |  | 18.5 | 0.051 | 0.93 | 0.047 | 0.94 |
| Condition 3 | 476877RR L4 TOM+ | L4 | 10826 | 342 | 10005 | 168 |  | 58.6 | 0.017 | 0.91 | 0.032 | 2.04 |
|  |  | L4 | 2474 | 61 | 1730 | 23 |  | 74.2 | 0.013 | 0.69 | 0.025 | 2.65 |
| Condition 3_o | 476877RR L4 TOM+ | L4 | 5845 | 100 | 5704 | 192 |  | 28.7 | 0.034 | 0.94 | 0.017 | 0.52 |
| Condition 4_o | 476877RL L4 TOM+ | L4 | 5877 | 244 | 5803 | 237 |  | 23.5 | 0.041 | 0.95 | 0.042 | 1.03 |
|  | **TOTAL** | **L4** | **63456** | **2274** | **58214** | **1910** | **AVE** | **34.2** | **0.0351** | **0.88** | **0.036** | **1.32** |
|  | stdev |  | 2996 | 119 | 2710 | 69 | stdev | 19.4 | 0.0145 | 0.08 | 0.013 | 0.68 |
| Condition 6_2_o | 487664RL P19 TOM+ | P19 | 2297 | 213 | 2179 | 224 |  | 8.7 | 0.103 | 0.85 | 0.093 | 0.95 |
| Condition 11 | 513583R P19 TOM+ | P19 | 10073 | 717 | 8271 | 855 |  | 8.7 | 0.103 | 0.74 | 0.071 | 0.84 |
|  |  | P19 | 10454 | 567 | 9122 | 429 |  | 20.3 | 0.047 | 0.83 | 0.054 | 1.32 |
| Condition 13 | 513583L P19 TOM+ | P19 | 16504 | 1278 | 15160 | 1262 |  | 11 | 0.083 | 0.84 | 0.077 | 1.01 |
|  |  | P19 | 16948 | 945 | 14552 | 1176 |  | 11.4 | 0.081 | 0.79 | 0.056 | 0.8 |
| Condition 15 | 529059L P19 TOM+ | P19 | 8031 | 778 | 7130 | 854 |  | 7.3 | 0.12 | 0.78 | 0.097 | 0.91 |
|  |  | P19 | 12101 | 696 | 10751 | 708 |  | 14.2 | 0.066 | 0.83 | 0.058 | 0.98 |
|  | **TOTAL** | **P19** | **76408** | **5194** | **67165** | **5508** | **AVE** | **11.7** | **0.086** | **0.81** | **0.072** | **0.97** |
|  | stdev |  | 5042 | 327 | 4471 | 374 | stdev | 4.4 | 0.025 | 0.04 | 0.02 | 0.17 |
|  | **2 tailed ttest** |  | **0.0328** | **0.0003** | **0.0461** | **0.0002** |  | **0.009** | **0.0001** | **0.04** | **0.0002** | **0.211** |
|  | P19 to L4 |  | * | **** | * | **** |  | * | ** | * | **** |  |

Notes: Each condition is an individual mouse. Each line within a condition is a technical replicate consisting of imaging from a separate piece of mammary tissue. L4 litters were not standardized, nor cross-fostered. Condition X\_o was imaged using tdTomato fluorescence, co-stained with RedDot2, and cleared using the CUBIC protocol. Others were re-stained with anti-SMA, co-stained with RedDot2, and cleared with a modified iDISCO protocol.

## Discussion

In the work presented here, we developed a modular hybrid analytical strategy combining deep-learning segmentation with classical image processing to segment and quantify alveoli, nuclei, and myoepithelial cells across intact mammary gland tissue volumes containing hundreds of thousands of cells. To evaluate this approach, we analyzed mammary gland tissue immediately before and after a well-characterized round of postpartum endoreplication in the mouse.^22–24^ This nuclear division cycle is thought to occur primarily in luminal epithelial cells and is proposed to increase secretory capacity during the onset of lactation. Using the combined MyoNet-AlveoliNet-Cellpose pipeline, we observed an approximately two-fold reduction in the fractional proportion of alveolar myoepithelial nuclei, from 8.6 ± 2.5% at pregnancy day 19 to 3.5 ± 1.4% at lactation day 4 (**Table 2**). These results are consistent with expansion of the luminal epithelial compartment during early lactation through endoreplication. In addition to the expected reduction in the proportion of myoepithelial nuclei, increased variability was observed among lactation day 4 samples. This variability may reflect localized autocrine or paracrine signaling mechanisms regulating proliferation, nuclear division, or endoreplication within individual alveoli during the early stages of lactation.

Myoepithelial cells play critical roles in lactation but have historically been difficult to study quantitatively. In conventional two-dimensional histological sections, only a small portion of an individual myoepithelial cell is typically visible because of its flattened morphology surrounding the alveolus. Furthermore, myoepithelial cells constitute only a small fraction of the total cellular population in the lactating gland. These characteristics make quantitative analysis using traditional approaches difficult and labor intensive. For example, manual scoring of individual alveoli in 60 µm tissue sections yielded approximately 150 alveolar epithelial cells per alveolus, with 7–11% classified as myoepithelial cells, which was insufficient for robust statistical analysis (data not shown). Similarly, scoring of 50 µm 3D sections using virtual reality visualization in syGlass modestly increased counts to approximately 200–300 cells per section but still required substantial manual effort by trained histologists and sampled only a small fraction of the tissue volume (data not shown).

Deep-learning–based segmentation provides several advantages over classical image segmentation methods, including tolerance to variations in signal intensity, robustness to differences in signal-to-noise ratio, reduced sensitivity to optical aberrations, and resilience to moderate morphological variability. These properties are particularly important for large volumetric biological datasets that may vary due to differences in tissue dissection, staining efficiency, reagent quality, tissue clearing, labeling intensity, or imaging depth. As microscopy datasets continue to increase in size and complexity, robust automated analysis methods become increasingly necessary.

For nuclear segmentation, recent advances in instance segmentation have enabled reliable quantification of nuclei in large 3D image volumes. In particular, StarDist and Cellpose have emerged as widely used and generalizable segmentation frameworks. After evaluating both approaches, we selected Cellpose because it provided more consistent segmentation performance at greater imaging depths in our datasets. The myoepithelial segmentation problem, however, required development of a specialized semantic segmentation model. To address this, we trained a custom 3D U-Net architecture using manually segmented volumetric datasets. The network was subsequently fine-tuned using multiple developmental stages, labeling strategies, and sample preparation methods to improve generalizability across experimental conditions. The U-Net architecture was particularly well suited for this application because it integrates broad volumetric tissue context with preservation of fine local boundaries through multiscale skip connections, enabling accurate segmentation of thin and spatially complex myoepithelial processes. The resulting MyoNet model produced robust segmentation across multiple experimental conditions and labeling modalities.

Despite the strong performance of MyoNet, false-positive segmentation occasionally occurred in structures exhibiting labeling patterns similar to myoepithelial cells. False positives were most commonly observed in ducts and vascular structures in tissues immunostained for smooth muscle actin (SMA). tdTomato expression was unexpectedly observed in a subset of stromal adipose cells.^31^ To address this limitation, we developed AlveoliNet to define the biological compartment for downstream quantitative analysis. Restricting quantification to nuclei located within segmented alveoli substantially improved specificity of myoepithelial cell identification. Without AlveoliNet filtering, myoepithelial counts were overestimated by approximately 25% in the OXTR-PA-tdTomato model and by approximately two-fold in SMA-immunostained tissues, with the largest effects observed at lactation day 4.

A major advantage of this modular analytical strategy is that semantic segmentation networks defining biological compartments can be combined with instance segmentation workflows to perform object-level quantification within anatomically constrained regions. This hybrid approach enabled quantitative identification of alveolar myoepithelial nuclei throughout intact tissue volumes while substantially reducing biologically implausible classifications. More broadly, this strategy demonstrates how compartment-aware deep-learning segmentation can be integrated with classical image analysis to study complex tissue organization in three dimensions.

All datasets presented here were acquired using spinning-disk confocal microscopy with a 40x 1.3 NA objective and 0.3 µm z-spacing. The segmentation workflow demonstrated consistent performance across multiple tissue clearing methods, labeling strategies, and imaging conditions, including tdTomato reporter expression and SMA immunostaining. Preliminary experiments further demonstrated compatibility with 3D time-lapse imaging of oxytocin-stimulated mammary explant contraction (**Movie 3**) suggesting potential utility for dynamic imaging studies. Future work will evaluate robustness across additional imaging modalities, magnifications, and sampling densities, as well as compatibility with additional myoepithelial markers including Mylk, Cnn1, and Krt5.

We initially developed MyoNet and AlveoliNet to study genetically modified mouse models exhibiting delayed lactation onset and apparent failure of normal postpartum endoreplication (Monks, in preparation). However, we anticipate that this approach will also be broadly useful for analysis of other mouse models of lactation insufficiency induced by diet, genetics, or pharmacological perturbation. Because core needle biopsy produces a relatively intact cylindrical tissue sample, we also foresee potential applications in the quantitative analysis of biopsy material from dairy species and humans.

Although initial MyoNet training was performed using the Alpine supercomputing system at the University of Colorado Boulder, subsequent optimized training strategies using smaller image windows and increased subsampling enabled effective model training on high-end desktop workstations equipped with modern GPUs. This substantially lowers the computational barrier for similar volumetric biological segmentation workflows.

In general, these studies demonstrate that integration of tissue clearing, volumetric imaging, deep-learning segmentation, and object-level analysis enables quantitative characterization of complex biological structures within intact tissues. We anticipate that similar modular 3D segmentation pipelines will be broadly applicable to studies of organ development, disease progression, drug response, and dynamic tissue remodeling across a wide range of biological systems.

## Supplemental Files

**Movie 1:** Myo Segmenting Movie.mp4

**Movie 2:** Condition_1_1-AlveoliNet&MyoNet.mp4

**Movie 3:** 4D MyoNet Segment.mp4

## Protocol Method: Using AlveoliNet and MyoNet

Detailed instructions for analysis of 3D image files implementing AlveoliNet and MyoNet.

## Data availability

The original Raw image files listed in Table 2 are available on FigShare (https://figshare.com/s/518e4fd1a86d21801117) SlideBook (.sldy) files are available and can be opened in SlideBook, SlideBook Reader (free), FIJI, or any program supporting Bioformats. U-Net kernals for both AlveoliNet and MyoNet are available as .pth files on FigShare. (https://figshare.com/s/518e4fd1a86d21801117)

The final masked image files resulting from the MyoNet-AlveoliNet-Cellpose pipeline for all data sets are archived here: https://monkslab.org/portal/ Movies displaying the results of the MyoNet-AlveoliNet-Cellpose pipeline for each are also available there.

## Lead contact

Further information for resources and reagents should be directed to and will be fulfilled by the lead contact, Jenifer Monks

## Materials availability

No unique/stable reagents were generated in this study.

## Software used

CellPose 3 syGlass (Morgantown, WV, USA)

SlideBook 2025 (Intelligent Imaging Innovations, Inc., Denver, CO)

CellNet 2.0 (Intelligent Imaging Innovations, Inc., Denver, CO)

GraphPad Prism 11 (DotMatics, Boston, MA)

## Author contributions

Colin Monks and Jenifer Monks contributed study conception and design. Creation and donation of the mouse strain was done by Yukiko U. Inoue and Takayoshi Inoue. ^.^Biological material preparation was performed by Kiarra Coger and Jenifer Monks. Data collection was performed by Colin Monks and Jenifer Monks. Analyses were performed by Cassidy Nicks, Sarah Albers, Colin Monks and Jenifer Monks. Initial MyoNet design, training and data hand segmentation was performed by Sarah Albers. The first draft of the manuscript was written by Colin Monks and Jenifer Monks. All authors read and approved the final manuscript.

Supplemental File: Protocol Method: Using AlveoliNet and MyoNet

First, collect multi-channel, 3-dimensional datasets using SlideBook software running Synergy. Synergy is a commercial module of SlideBook that allows external applications running the Synergy protocol to share data directly.

Next, apply neural networks to the data to generate binary masks of 3D structures. We load the AlveoliNet weights (AlveoliNet.pth) into CellNet 2.0 (Intelligent Imaging Innovations, Inc.), a Synergy application for generating and applying deep learning neural networks. CellNet 2.0 can run on any Windows 11 computer running SlideBook 2025 or later with the Synergy Module. CellNet 2.0 supports CPU, GPU (CUDA), and OpenVino environments. High-powered, modern NVIDIA graphics cards support faster computations and larger datasets. We apply AlveoliNet to the RedDot2 channel using a subsampling factor of 0.1 to match the data to the network geometry and a probability threshold cutoff of 0.8. Although the threshold variable is relatively insensitive, keep it high to reduce false positive identification of alveolar structures. CellNet 2.0 automatically returns the AlveoliNet result as a 3D binary mask associated with the original data in SlideBook.

Then repeat the process with MyoNet (MyoNet.pth) loaded into CellNet 2.0. Apply MyoNet to the tdTomato or SMA channel using a subsampling factor of 0.25 and a probability cutoff of 0.5 to match the network geometry and suppress false positives. CellNet 2.0 automatically returns the resulting mask to SlideBook.

Next, use CellNet 2.0 to apply CellPose3 with the ‘nuclei’ weight set to the RedDot2 data using a 10 µm nuclei size setting. This process returns an instance segmentation in which the system assigns a unique identifier to each nucleus within the 3D SlideBook mask.

After generating all masks (AlveoliNet, MyoNet, and CellPose nuclei), refine them using binary AND mask operations that preserve instance segmentation. Perform these operations using the binary mask refinement functions in Synergy Conductor (Intelligent Imaging Innovations, Inc.). We identify alveolar nuclei by preserving any nucleus instance that overlaps with the AlveoliNet mask by at least 80%. From this subset, we identify myoepithelial nuclei as those alveolar nuclei that overlap with the MyoNet mask by at least 50%.

We select this gating strategy to balance sensitivity and specificity, particularly to reduce false negatives in SMA-labeled myoepithelial cells, where staining typically does not fully cover the nuclear region. In contrast, tdTomato labeling provides robust nuclear coverage and remains insensitive to this gating threshold.

This mask refinement procedure produces instance-segmented populations of total nuclei, alveolar nuclei, and myoepithelial nuclei. The system automatically pushes each resulting mask to the original SlideBook dataset as a 3D mask, with instance counts precomputed and recorded in the mask name.

Evaluate masks in each data set by empirical simple visual evaluation prior to proceeding to the next step. Enter mask instance counts into GraphPad Prism for statistical analysis.

## Acknowledgments

We would like to dedicate this work to the memory of Dr. Abraham “Avi” Kupfer who inspired our deep interest in quantitative microscopy. We thank Dr. Dan Krofcheck (University of New Mexico) for his helpful discussion. This work was supported by the Alpine HPC system, which is jointly funded by the University of Colorado Boulder, the University of Colorado Anschutz, Colorado State University, and the National Science Foundation (award 2201538). Imaging time and resources were contributed by Intelligent Imaging Innovations, Inc. (3i). Additional resources were provided by University of Colorado School of Medicine Bridge funding (Monks, J). Research reported in this publication was supported by the Eunice Kennedy Shriver National Institute of Child Health & Human Development of the National Institutes of Health under Award Number R01HD117769 (Monks, J). The content is solely the responsibility of the authors and does not necessarily represent the official views of the National Institutes of Health.

## References

1 Medina, D. Preneoplastic lesions in murine mammary cancer. Cancer Res 36, 2589–2595 (1976).

2 Tolg, C., Cowman, M. & Turley, E. A. Mouse Mammary Gland Whole Mount Preparation and Analysis. Bio Protoc 8, e2915 (2018). 10.21769/BioProtoc.2915

3 Brisken, C. & Ataca, D. Endocrine hormones and local signals during the development of the mouse mammary gland. Wiley Interdiscip Rev Dev Biol 4, 181–195 (2015). 10.1002/wdev.172

4 Hennighausen, L. & Robinson, G. W. Information networks in the mammary gland. Nat Rev Mol Cell Biol 6, 715–725 (2005). 10.1038/nrm1714

5 Rooney, B. L., Rooney, B. P., Muralidaran, V., Wang, W. & Furth, P. A. Mouse Mammary Gland Whole Mount Density Assessment across Different Morphologies Using a Bifurcated Program for Image Processing. Am J Pathol 192, 1407–1417 (2022). 10.1016/j.ajpath.2022.06.013

6 Orlicky, D. J. et al. Using the novel pelvic organ prolapse histologic quantification system to identify phenotypes in uterosacral ligaments in women with pelvic organ prolapse. Am J Obstet Gynecol 224, 67 e61–67 e18 (2021). 10.1016/j.ajog.2020.10.040

7 Ogony, J. et al. Towards defining morphologic parameters of normal parous and nulliparous breast tissues by artificial intelligence. Breast Cancer Res 24, 45 (2022). 10.1186/s13058-022-01541-z

8 Dawson, C. A. & Visvader, J. E. The Cellular Organization of the Mammary Gland: Insights From Microscopy. J Mammary Gland Biol Neoplasia 26, 71–85 (2021). 10.1007/s10911-021-09483-6

9 Davis, F. M. et al. Single-cell lineage tracing in the mammary gland reveals stochastic clonal dispersion of stem/progenitor cell progeny. Nat Commun 7, 13053 (2016). 10.1038/ncomms13053

10 Stewart, T. A., Hughes, K., Hume, D. A. & Davis, F. M. Developmental Stage-Specific Distribution of Macrophages in Mouse Mammary Gland. Front Cell Dev Biol 7, 250 (2019). 10.3389/fcell.2019.00250

11 Hitchcock, J. R., Hughes, K., Harris, O. B. & Watson, C. J. Dynamic architectural interplay between leucocytes and mammary epithelial cells. FEBS J 287, 250–266 (2020). 10.1111/febs.15126

12 Dawson, C. A. et al. Tissue-resident ductal macrophages survey the mammary epithelium and facilitate tissue remodelling. Nat Cell Biol 22, 546–558 (2020). 10.1038/s41556-020-0505-0

13 Corral, D. et al. Mammary intraepithelial lymphocytes promote lactogenesis and offspring fitness. Cell 188, 1662–1680 e1624 (2025). 10.1016/j.cell.2025.01.028

14 Schmidt, U. W., M.; Broaddus, C.; Myers, G. in *Medical Image Computing and Computer Assisted Intervention – MICCAI 2018: 21st International Conference.* (ed A.F.; Schnabel Frangi, J.A.; Davatzikos, C.; Alberola-López, C.; Fichtinger, G.) (Springer-Verlag).

15 Weigert, M. S., U.; Haase, R.; Sugawara, K.; Myers, G. in 2020 IEEE Winter Conference on Applications of Computer Vision (WACV).

16 Weigert, M. S., U. . in 2022 IEEE International Symposium on Biomedical Imaging Challenges (ISBIC). (IEEE).

17 Pachitariu, M. & Stringer, C. Cellpose 2.0: how to train your own model. Nat Methods 19, 1634–1641 (2022). 10.1038/s41592-022-01663-4

18 Falk, T. et al. U-Net: deep learning for cell counting, detection, and morphometry. Nat Methods 16, 67–70 (2019). 10.1038/s41592-018-0261-2

19 Serra, J. Image Analysis and Mathematical Morphology. Vol. 1 (Academic Press, Inc., 1982).

20 Inoue, Y. U. et al. Targeting Neurons with Functional Oxytocin Receptors: A Novel Set of Simple Knock-In Mouse Lines for Oxytocin Receptor Visualization and Manipulation. eNeuro 9 (2022). 10.1523/ENEURO.0423-21.2022

21 Renier, N. et al. iDISCO: a simple, rapid method to immunolabel large tissue samples for volume imaging. Cell 159, 896–910 (2014). 10.1016/j.cell.2014.10.010

22 Linkert, M. et al. Metadata matters: access to image data in the real world. J Cell Biol 189, 777–782 (2010). 10.1083/jcb.201004104

23 Virtanen, P. et al. SciPy 1.0: fundamental algorithms for scientific computing in Python. Nat Methods 17, 261–272 (2020). 10.1038/s41592-019-0686-2

24. patchify (version 0.2.3) v. 0.2.3 (2017).

25 Solovyev, R., Kalinin, A. A. & Gabruseva, T. 3D convolutional neural networks for stalled brain capillary detection. Comput Biol Med 141, 105089 (2022). 10.1016/j.compbiomed.2021.105089

26 Kingma, D. P. J. B., J. in 3rd International Conference for Learning Representations. (Arxiv).

27 Salehi, S. S. M. E., D.; Gholipour, A. Tversky loss function for image segmentation using 3D fully convolutional deep networks. (2017). 10.1007/978-3-319-67389-9_44

28 Paszke, A. G., S.; Massa, F.; Lerer, A.; Bradbury, J.; Chanan, G.; Killeen, T.; Lin, Z.; Gimelshein, N.; Antiga, L.; Desmaison, A.; Köpf, A.; Yang, E.; DeVito, Z.; Raison, M.; Tejani, A.; Chilamkurthy, S.; Steiner, B.; Fang, L.; Bai, J.; Chintala, S. PyTorch: An Imperative Style, High-Performance Deep Learning Library. Arxiv (2019).

29 Abadi, M. B., P.; Chen, J.; Chen, Z.; Davis, A.; Dean, J.; Devin, M.; Ghemawat, S.; Irving, G.; Isard, M.; Kudlur, M.; Levenberg, J.; Monga, R.; Moore, S.; Murray, D.G.; Steiner, B.; Tucker, P.; Vasudevan, V.; Warden, P.; Wicke, M.; Yu, Y.; Zheng, X.;. in 12th USENIX Symposium on Operating Systems Design and Implementation (OSDI ’16).

30 Open Neural Network Exchange (ONNX) v. v1.21.0 (2017).

31 Li, E. et al. Oxytocin signaling in adipocytes is required for normal milk fat production. Cell Metab (2026). 10.1016/j.cmet.2026.03.013

